# Pollinator preference for large flowers persists when climate induces the expression of smaller flowers

**DOI:** 10.64898/2026.10.12.758611

**Authors:** Samantha Day Briggs, Mallory C. Howard, Jill T. Anderson

## Abstract

**Premise:** In arid environments, large flowers can be disadvantageous owing to excess transpiration through petals and other floral organs, often resulting in the expression of smaller flowers. Larger flowers, which serve as visual attractants, can provide greater rewards compared to smaller flowers resulting in a greater number of floral visitors. Environmental conditions can also shift community compositions of pollinators across landscapes resulting in different preferences. Therefore, pollinator preferences and abiotic factors may conflict or act synergistically to shape floral size across elevation.

**Methods:** For four years, we examined how *Iris missouriensis* (Iridaceae) flower size changes across dynamic abiotic and biotic conditions. We monitored populations along a steep environmental gradient in the Colorado Rocky Mountains to investigate the relationship of flower size, elevation, and climate. For two of the years, we also established arrays of artificial flowers across the same gradient to evaluate the flower size preference of floral visitors, while controlling for floral traits other than size. We recorded visitations to these arrays across two years to capture interannual variation in the preferences of floral visitors.

**Results:** Consistent with expectations based on clinal gradients in precipitation, we found that flower size increased with elevation in most years, resulting in smaller flowers in arid low elevation sites. However, that pattern was lost in 2023 when temperatures were the coolest and early season precipitation was high. In our artificial floral arrays, larger flowers received more visitors relative to smaller flowers even in sites where naturally recruiting plants had much smaller flowers. These results are consistent with climate-induced clinal variation in *I. missouriensis* floral size where floral visitor preference does not track local flower size.

**Conclusions:** As climate change progresses, conflicting effects from pollinators and abiotic forces could alter the expression and evolution of flower size, potentially disrupting plant reproduction for animal-pollinated species. Consistent pollinator preference for large flowers, even in sites with naturally small flowers, indicates a potential mismatch between biotic and abiotic forces on floral trait expression.

## Introduction

Floral sizes vary dramatically across taxa, with the smallest flower in the world at roughly 0.5 mm in diameter, of the genus *Wolffia*, requiring a microscope for observation (Schmitz & Kelm, 2017), and the largest flower, belonging to the genus *Rafflesia*, growing to around one meter in diameter (Wicaksono et al., 2016). This wide diversity in floral displays is often attributed to selection driven by pollinator preferences (Caruso et al., 2019). Indeed, pollinator-mediated selection can act rapidly to increase flower size across generations (Ramos & Schiestl, 2019). Diverse pollinator species often prefer large flowers, likely due to their increased nectar and pollen rewards relative to smaller flowers (Ortiz et al., 2021). Large flowers can also attract a greater number of floral visitors because they can provide shelter and heat as a reward for small pollinating insects (Lozada-Gobilard, Nielsen, et al., 2023). This pattern is the basis of the pollinator market framework which suggests that pollinators choose which flowers to visit based on the ‘advertisements’ of their rewards (Filella et al., 2013).

In addition to pollinators, other biotic interactions (Parachnowitsch & Caruso, 2008; Sletvold, 2019; Strauss & Whittall, 2006) and abiotic drivers such as drought (Descamps et al., 2021) and elevated temperatures (Alquichire-Rojas et al., 2024) influence these floral traits. Abiotic factors can reinforce or even counteract the effects of pollinators (Caruso et al., 2019; Day Briggs & Anderson, 2025; Kuppler et al., 2021; Sletvold, 2019). For example, in *Ipomoea purpurea* (Convolvulaceae), pollinators favor larger flowers, but drought induces the production of smaller flowers (García et al., 2023), strengthening the effect of pollinator preference under drought conditions. When evaluating global patterns of natural selection in plants, up to 67% of the variation can be explained by patterns of precipitation (Siepielski et al., 2017). Indeed, drought stress is a strong driver of trait expression and evolution that can reduce traits associated with pollinator attraction (Phillips et al., 2018) and can increase the cost of maintaining flowers, leading to the expression of smaller flowers (Galen, 2000; Gallagher & Campbell, 2017; Roddy et al., 2023). In animal-pollinated flowering plants, the expression of flower size could be driven by a tradeoff between pollinator preference for larger flowers and the expression of flower size as a result of moisture availability (Irwin et al., 2001; Krizek & Anderson, 2013; Navarro & Medel, 2009; Parachnowitsch & Caruso, 2008; Sletvold et al., 2015).

Although pollinators often prefer larger floral displays (Chapurlat et al., 2015; Knauer & Schiestl, 2017; Lavi & Sapir, 2015), the strength of these preferences varies depending on community compositions. Pollinator community compositions can shift across environmental gradients in response to abiotic conditions (Lefebvre et al., 2018; McCabe et al., 2022; Ogilvie & CaraDonna, 2022; WallisDeVries et al., 2011). For example, butterfly abundance decreases significantly under drought conditions (WallisDeVries et al., 2011), and flies are more abundant at high elevations than other pollinators (McCabe et al., 2022). Functional groups of pollinators often differ in their floral preferences (Herrera, 2020; Thompson, 2001), as is the basis for pollination syndromes (Dellinger, 2020). Thus, spatial variation in pollinator community assemblages can result in floral trait preferences that shift across the landscape (Gómez et al., 2008), potentially interacting with the direct effects of climate, which could generate clinal variation in floral traits. However, most studies of pollinator preference for floral traits are conducted at a single site or under controlled conditions, so it remains unclear whether preferences track local variation in floral traits when both abiotic conditions and biotic communities are shifting across the landscape.

Floral traits associated with pollinator attraction, such as flower size, typically serve as honest cues of floral rewards such as nectar and pollen (Blarer et al., 2002; Fenster et al., 2006; Knauer & Schiestl, 2015; Lozada-Gobilard, Nielsen, et al., 2023). Correlations of floral rewards and floral size can make it challenging to estimate pollinator preference for flower size alone. Researchers have employed artificial flowers to determine that pollinator preference for floral shapes is evolved rather than learned (Howard et al., 2019), and that larger UV-absorbing spots reduce the amount of UV light reflected onto anthers (Koski & Ashman, 2015). The use of artificial flowers in studies such as these enable researchers to generate models with trait variation that goes beyond what is found in natural populations, while controlling for other traits that co-vary with morphology, such as scent, nectar and pollen availability, or color patterns.

We examined how flower size of the Rocky Mountain Iris (*Iris missouriensis* Nutt., Iridaceae) varies across an elevational gradient where aridity and pollinator community composition (Day Briggs & Anderson, 2026) covary. Species of the Iridaceae generally have large flowers that attract diverse pollinators and they often occur in arid environments (Angiolini et al., 2015; Arafeh et al., 2002; Lavi & Sapir, 2015), making them well suited for examining whether aridity and pollinator preferences conflict or act synergistically in their potential influences on the expression of floral traits. To test whether flower size covaries with aridity, we measured flower size of naturally recruiting *I. missouriensis* plants in multiple populations across a broad elevational range for four years, allowing us to assess whether clinal patterns were consistent across years that differed in temperature and precipitation. To determine how the preference of floral visitors varies across elevation and control for variation in other traits that co-vary with size, such as rewards or other attractants, we performed observation studies with 3D printed artificial flowers. Using these floral models, we then tested whether floral visitors of *I. missouriensis* (i) preferentially visit the largest flowers available at all sites across the elevational gradient, or (ii) preferentially visit flowers of the average size at each site across the gradient. Together, these approaches allowed us to evaluate whether floral visitor preferences track or oppose the aridity-associated patterns in flower size along a gradient where both abiotic conditions and visitor communities change.

## Methods

*Iris missouriensis* is a perennial forb that is pollinated by a diversity of animals including bees, butterflies, hummingbirds, ants, and flies, with community compositions that shift across elevation (Day Briggs & Anderson, 2026). It has large, light purple flowers that produce nectar and provide shelter for small insects (Fig. 1). This species inhabits wet meadows in the western half of North America, ranging from Northern Mexico to Alaska (Cornman & Arnold, 2007; Day Briggs & Anderson, 2026).

**Figure 1:**
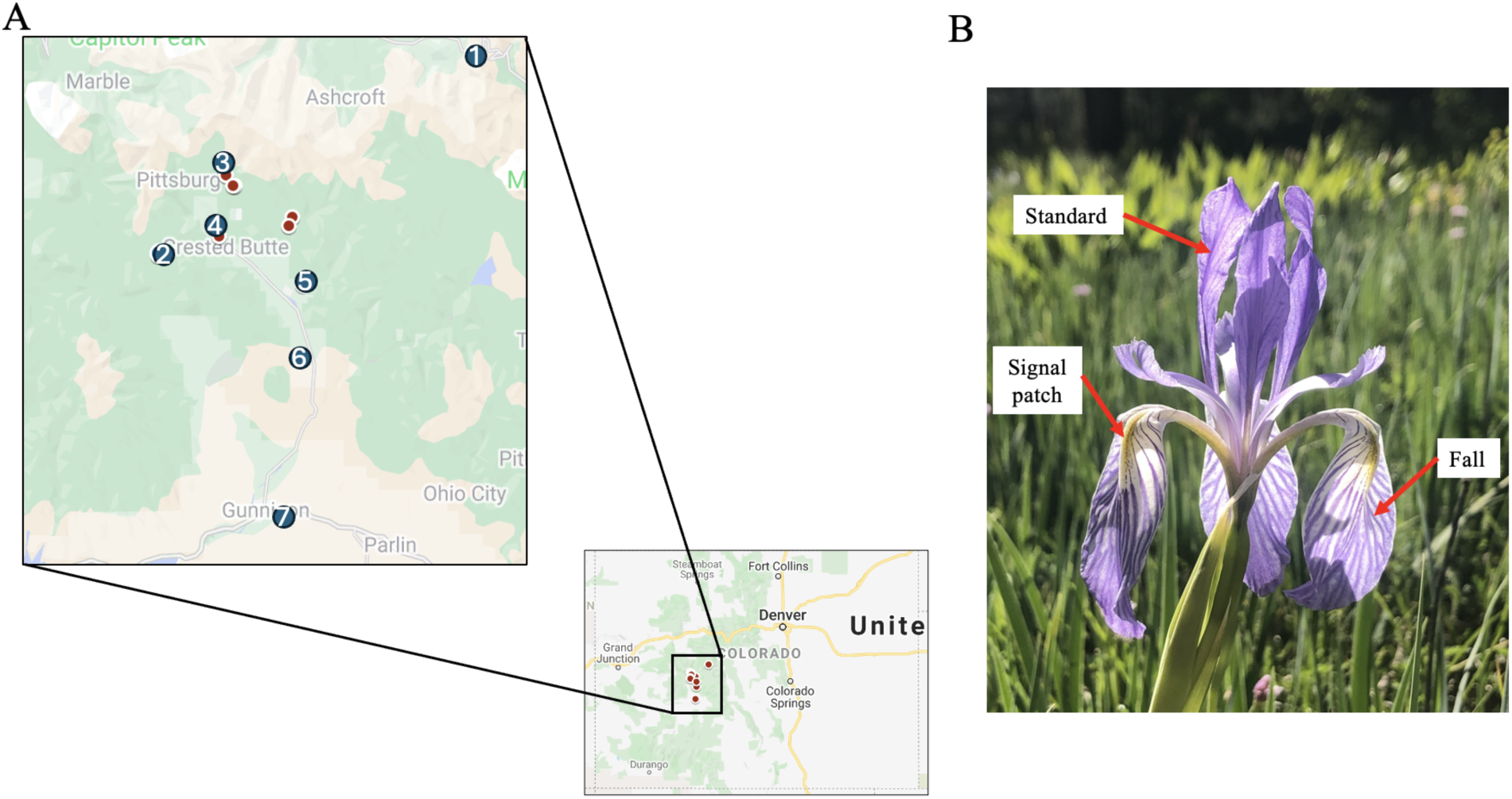
Study sites and study system. A) Indicates all of the study sites where natural flowers were measured (blue and red circles) and the sites where artificial floral arrays were set out and recorded (blue numbered circles only). B) Demonstrates the *Iris missouriensis* floral shape and morphology. The fall and standard tepals are identified along with the signal patch.

We conducted fieldwork in the Elk Mountain Range of Gothic, Colorado (USA), near the Rocky Mountain Biological Laboratory from 2020 to 2023 (Fig. 1). In this region, higher elevations have more mesic climates compared with arid low elevation locations (Anderson & Wadgymar, 2020). This region has an abundant and diverse pollinator community that has been extensively cataloged (Forrest & Thomson, 2011; Inouye, 1978; McKinney et al., 2012; Syskine & Boggs, 2024).

### Clinal variation in flower size across natural populations

To assess phenotypic clines in flower size, we identified N= 16 natural populations of *I. missouriensis* (elevations: 2337-3331m; Fig. 1A) using verified records in GBIF.org, online herbariums, and research grade iNaturalist entries. Each growing season, during peak flowering, we visited all 16 populations and measured the length and width of the fall and standard tepal (a petaloid sepal), using a single set of digital calipers on up to 20 individuals per population (Fig. 1B). We multiplied length and width to calculate tepal area and added the area of falls and standards to generate the measurement of floral area (as in Sapir et al., 2002). *Iris missouriensis* grows clonally; therefore, we sampled individuals at least one meter apart to ensure that we measured genetically distinct individuals.

Higher elevation sites in this region generally have cooler temperatures and greater moisture availability (Anderson & Wadgymar, 2020; Dunne et al., 2003). To confirm this relationship between climate and elevation in our study, we extracted interpolated April, May, and June mean precipitation and June average temperatures for all sites during the study years 2020-2023 from the Parameter-elevation Relationships on Independent Slopes Model (PRISM) database (Daly et al., 2008). These populations tend to reach peak flowering throughout the month of June after snowmelt begins in April, so this timeframe represents the average moisture (as melting snowpack and growing season rain) and temperatures experienced by these plants during peak reproduction. We regressed climatic variables independently on elevation, year, and month for precipitation, along with their interaction, with a random effect of study site using the ‘lmer’ function in the the lme4 package v. 1.1-36 (Bates et al., 2015) in R v. 4.4.1 statistical software (R Core Team, 2024)

To test our hypothesis that floral size increases elevation, we conducted a linear mixed effects model using the ‘lmer’ function in the the lme4 package v. 1.1-36 (Bates et al., 2015) in R v. 4.4.1 statistical software (R Core Team, 2024) to evaluate flower size as a function of elevation, using year of data collection as a fixed effect to account for multiple years of data collection, and interactions of year and elevation. We also included the study site as a random effect in our model and used a normal distribution. We used the R package DHARMa to confirm that residuals were normally distributed and homoscedastic (Hartig, 2024).

Clinal variation in traits can also arise through stochastic processes related to isolation-by-distance (Lande, 1991). Thus, as a first attempt to capture genetic structure we conducted a preliminary, small-scale, study to assess gene flow between populations by sampling leaf tissue from 64 individuals from 15 of the monitored populations (N = 3-6 individuals per population). We initially identified eight microsatellite markers developed for congeneric *Iris* species that were effective for evaluating genetic diversity (Burke & Arnold, 1999; Cornman & Arnold, 2007; Meerow et al., 2005; Sun et al., 2012). Through this testing, we determined that the nuclear markers IEST12 and IEST24, and the chloroplast marker TrnL, were the best candidates for capturing population structure in these *Iris missouriensis* populations. Using the selected primers, we amplified DNA and sent samples for Sanger Sequencing at Eurofins Genomics (Louisville, KY). We used Geneious Prime (2024.0.7, https://www.geneious.com) to clean and trim sequences. We used the R package adegenet v. 2.1.11 (Jombart, 2008) to perform discriminate analysis of principal components (DAPC) allowing us to identify genetic clusters, then the package hierFstat v. 0.5-11 (Goudet, 2005) to calculate differentiation between populations as F_st_.

### Artificial floral arrays

We leveraged our measurements of flowers from natural populations in the 2020 field season to generate artificial flowers in nine size categories ranging from 2427.43 mm^2^ to 11625.32 mm^2^. This range was calculated based on the average size and four standard deviations above and below the size we measured across all natural populations (Fig. 2A), resulting in artificial flowers with greater variation in size than in any natural population. We created these three dimensional models using Tinkercad (Autodesk, 2021) and printed using an Ender 3 Creality 3D printer (2017, Creality.com) in white compostable filament (Fig. 2A). We manually painted the flowers to match colors and signal patterning of naturally occurring *I. missouriensis* (Fig. 2B). *Iris missouriensis* produces negligible ultraviolet pigment patterning when viewed under a UV light source (The Authors, unpublished finding).

**Figure 2:**
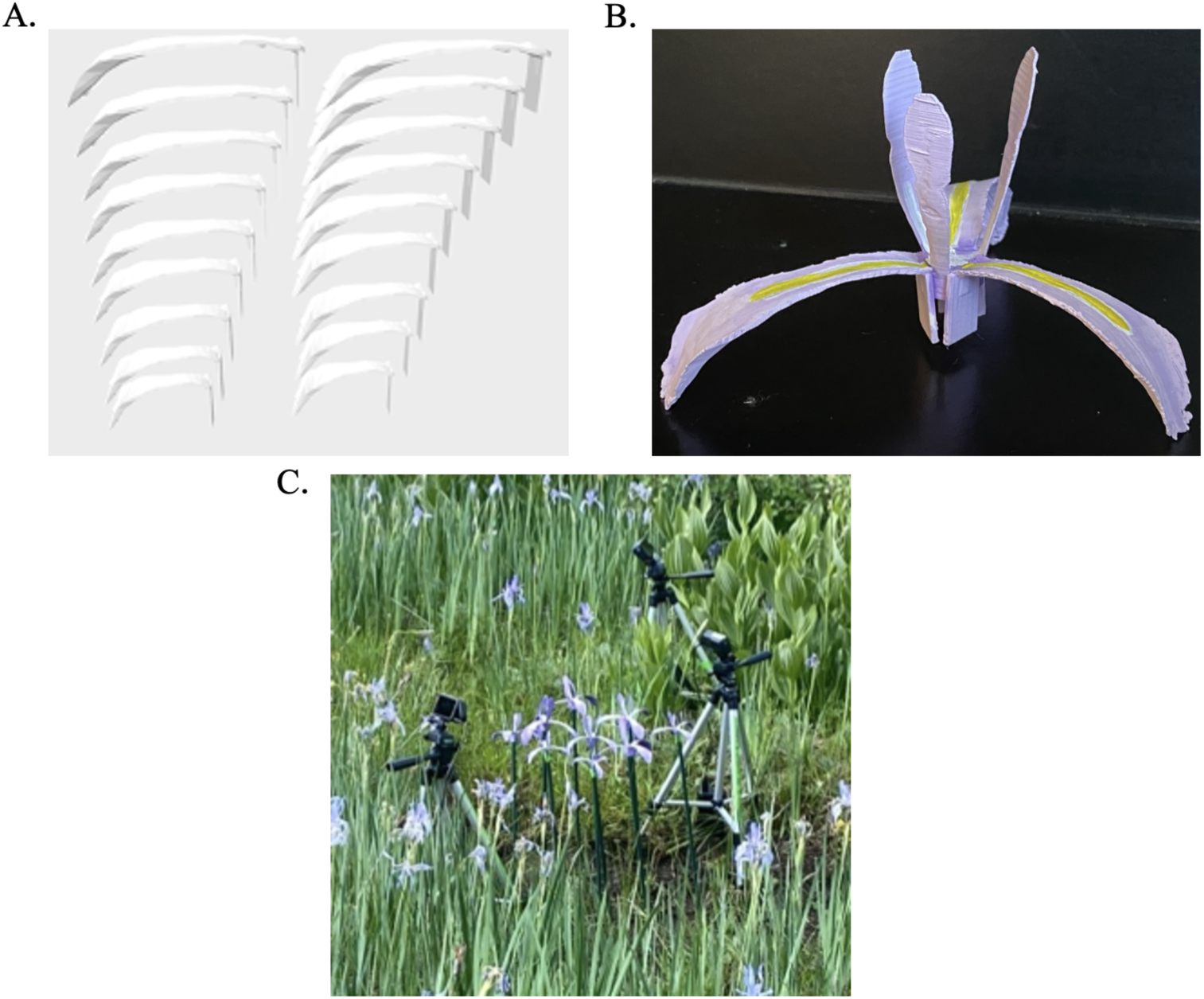
3D printed artificial flower design and array setup. A) A 2D image of the 3D .gcode file showing the tepals of all nine size categories. This is the orientation that the flowers were printed. B) Shows an image of an assembled and painted 3D printed *Iris missouriensis* flower. C) Depicts the artificial floral array setup with cameras surrounded by natural *I. missouriensis* flowers.

During the summers of 2021 and 2022, we fastened the nine 3D printed flowers of different sizes to 0.5 m tall metal rods and placed them in a haphazard order in a 3×3 floral array (Fig. 2C). We used GoPro Hero8 cameras (2020, GoPro Inc.) in full HD 1920×1080 pixel quality using 256 GB microSD cards to record floral visitations (Fig. 2C). A foundational study found no significant difference in pollinator data collected from recordings vs. in person observations (Gilpin et al., 2017). We recorded each array from two camera vantage points, in front and behind the grided array, to provide multiple angles of view when documenting floral visitors. We established arrays and recorded on clear days with little cloud cover to ensure peak pollinator activity (Konvicka et al., 2023; Lawson & Rands, 2019). We recorded for an average of 5 hours at each site, but some recordings were shortened by unexpected adverse weather conditions and some recordings ran slightly longer (range of recording lengths: 2 hours and 57 minutes to 8 hours and16 minutes). We set up these arrays within seven previously identified natural *I. missouriensis* populations across the elevational range (2337m, 2527m, 2695m, 2723m, 2937m, 3010m and 3331m; Fig. 2C). These recordings were performed at two time points about one week apart at each site during peak flowering, then repeated for a second season (2022) to account for temporal and interannual variation in visitor community (Table S1).

To determine whether our artificial flowers attracted similar assemblages of floral visitors as natural *I. missouriensis* flowers, we performed contemporaneous recordings on natural flowers at each site. We recorded visitations to 1-3 natural flowers using two GoPro Hero8 cameras (2020, GoPro Inc.) recording in 1920×1080 pixel quality with 256 GB microSD cards. To capture temporal variation in visitations, we recorded natural flower visitations for one hour in the morning, one hour at midday, and one hour in the evening.

### Floral visitor identification

To identify potential pollinators, we generated an insect guide based on the extensive pollinator research that has been conducted around the Rocky Mountain Biological Laboratory (e.g. Forrest & Thomson, 2011; Gezon et al., 2015; Stemkovski et al., 2020). We also used guides from The University of Colorado Boulder (Scott et al., 2011), iNaturalist (*iNaturalist*, 2024), and the Global Biodiversity Information Facility (GBIF, 2024) for identifications. We reviewed all recordings throughout the academic year with the help of five undergraduate research assistants. These assistants watched all video footage, documented the timestamps of each floral visitation, and identified visitors to broad categories. We only recorded an individual as a floral visitor if it made contact with the artificial or natural flower by touching or landing on the structure. The first author then revisited each recorded timestamp to identify floral visitors to at least family level. We submitted images from recordings to iNaturalist in the name of (The First Author) where experts contributed to the identification of 15 of the submitted floral visitors.

To determine if the artificial flowers attracted the same floral visitors as naturally-recruiting of *I. missouriensis* individuals, we rarified the dataset to a standard sample size to account for differences in sampling efforts using using the ‘rrarefy’ function in the vegan (v. 2.6-8) R package (Oksanen, 2010) in R v. 4.4.1 (R Core Team, 2024). We then generated a distance matrix of our floral visitor data using the ‘vegdist’ function in vegan with a Bray-Curtis index that we passed to the ‘adonis2’ function for permutational multivariate analysis of variance (PERMANOVA).

We retained all visitors for analyses, not just species traditionally classified as pollinators, because non-Syrphidae flies (Orford et al., 2015) and ants (Das & Das, 2023) are effective pollinators in some systems. However, we did conduct a secondary analysis with only bee, butterfly, and hummingbird visitors to determine the preferences of species considered effective pollinators across systems (Page et al., 2021). To examine relative visitation rates to artificial flowers in relation to the average size of natural flowers, we calculated the average floral size in the natural populations at each site in the two years of the study, then subtracted the average from the size of each artificial flower. We then modeled the number of visits as a function of the difference of flower size from average, we also included elevation and year of recording along with the interaction of size and year as fixed effects in our model. We also included a quadratic term for difference of flower size from average as an attempt to identify a possible limit to flower size preference. Due to overdispersion of the data, we used a generalized linear mixed effects model with a negative binomial distribution using the ‘glmmTMB’ function in the glmmTMB package (v. 1.1.10) (Brooks et al., 2017). We again used the R package DHARMa to confirm that residuals were normally distributed and homoscedastic (Hartig, 2024) and we plotted the results using the ‘ggpredict’ function of the ggeffects package v. 2.2.1 (Lüdecke, 2018).

## Results

### Clinal variation in flower size

Flower size varied significantly across elevation, though this pattern depended on year (elevation x year: *χ*² = 25.36, p < 0.0001, Fig. 3). We conducted post hoc tests to evaluate elevational trends for each year and found that floral area increased with elevation significantly in 2021 and 2022, but the trend was nonsignificant in 2020 and 2023. Pairwise contrasts indicate that the elevational response in 2023 differed significantly from 2021 and 2022 (Tukey-adjusted, p < 0.001) and approaching significance from 2020 (Tukey-adjusted, p = 0.076), but other comparisons were nonsignificant. Thus, we found a dynamic cline in flower size across elevation indicating that the environmental conditions that mediate this cline likely fluctuate across years.

**Figure 3:**
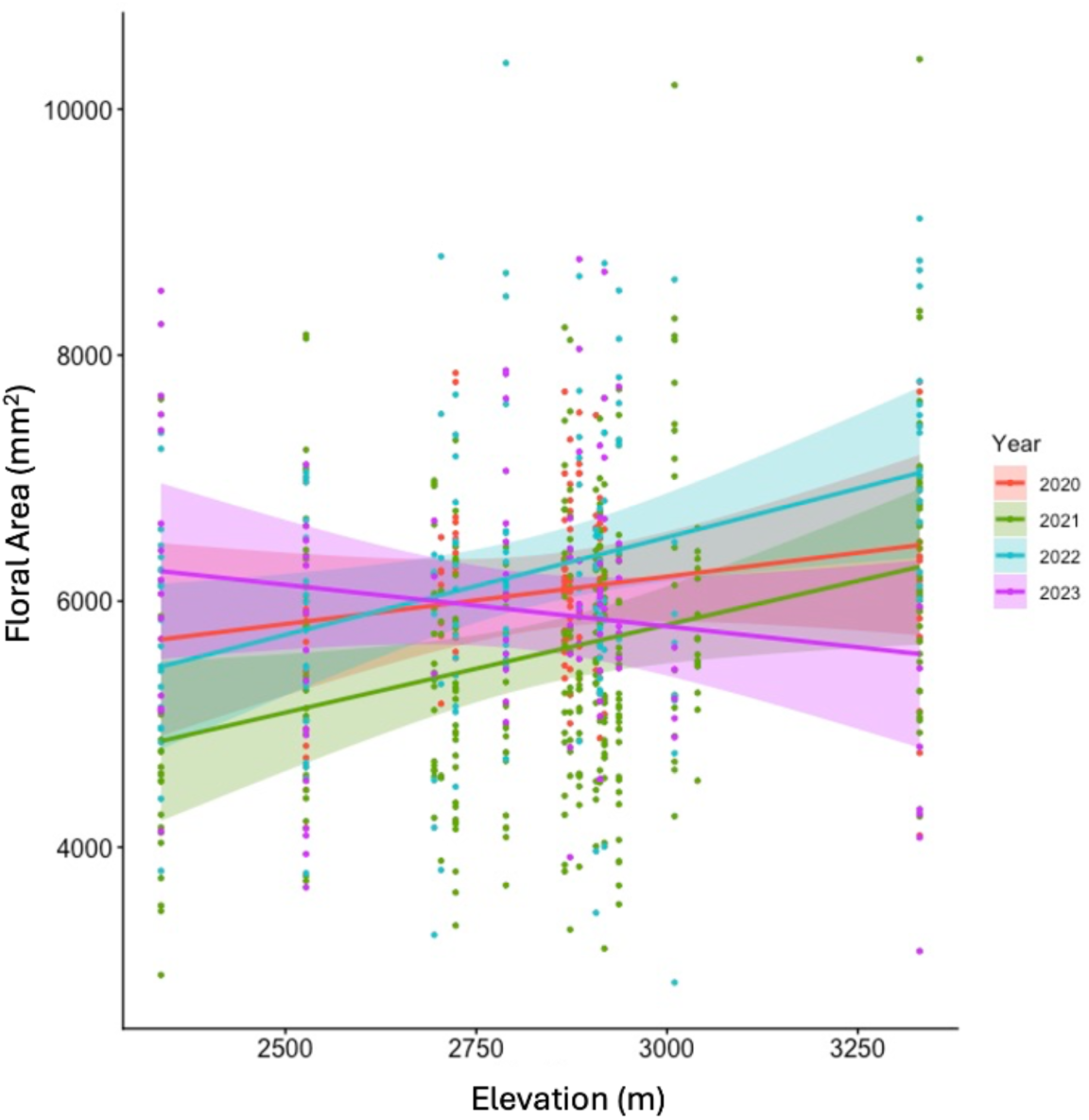
Dynamic cline in flower size across elevation. Flowers were larger at higher elevation sites, but this pattern depended on year.

Consistent with previous work (*i.e.* Anderson & Wadgymar, 2020; Dunne et al., 2003), we determined that climate — average growing season precipitation and June temperatures — were highly correlated with elevation across our sites. Precipitation increased with elevation, but depended on month and year (elevation x month x year: *χ*² = 44.41, p < 0.001, Fig. 4A) and that average June temperatures were cooler at higher elevations (*χ*² = 106.97, p < 0.001, Fig. 4B), but the strength of this relationship varied across years (elevation x year: *χ*² = 1401.04, p < 0.001, Fig. 4B). Pairwise comparisons of estimated marginal means indicate that temperatures of all years differed significantly from one another with 2023 being the coldest year (Tukey-adjusted p < 0.001 for all contrasts, Fig. 4B) and that April and June precipitation was the greatest in 2023 when averaged across elevations.

**Figure 4:**
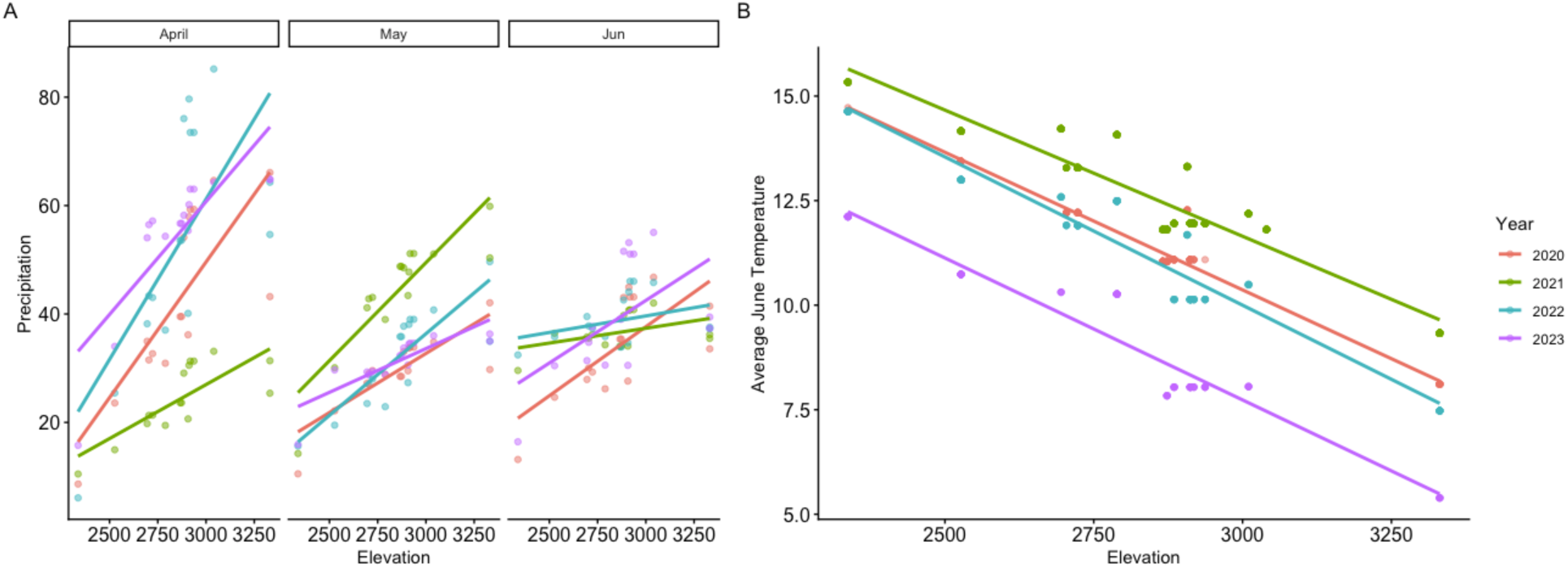
Average monthly precipitation as rain and melted snow for April, May, and June (A) and average June temperature (B) across sites and years of the study.

In addition, our small-scale genotyping study suggested high rates of gene flow between populations. Our microsatellite study indicated limited genetic differentiation between the 15 populations with a maximum of K = 5 supported (Fig. S1) and relatively low pairwise F_ST_ values in our nuclear markers (mean IEST12 F_ST_: 0.008, mean IEST24 F_ST_: 0.017, Fig. S2). There was insufficient variation in the chloroplast marker to calculate F_ST_.

### Floral visitor preference for flower size

We recorded a total of 2791 interactions with our artificial flowers, which attracted 37 different species of potential pollinators including bees, butterflies, and hummingbirds. The most common floral visitors were ants and flies (Fig. 5). The floral visitor communities of the natural and artificial flowers did not differ statistically (F_1,26_ = 1.48, p = 0.199, Fig. S3). Nevertheless, natural flowers hosted 33 unique species, artificial flowers had 11 unique species, and they shared visitations with 26 different species (Table S2).

**Figure 5:**
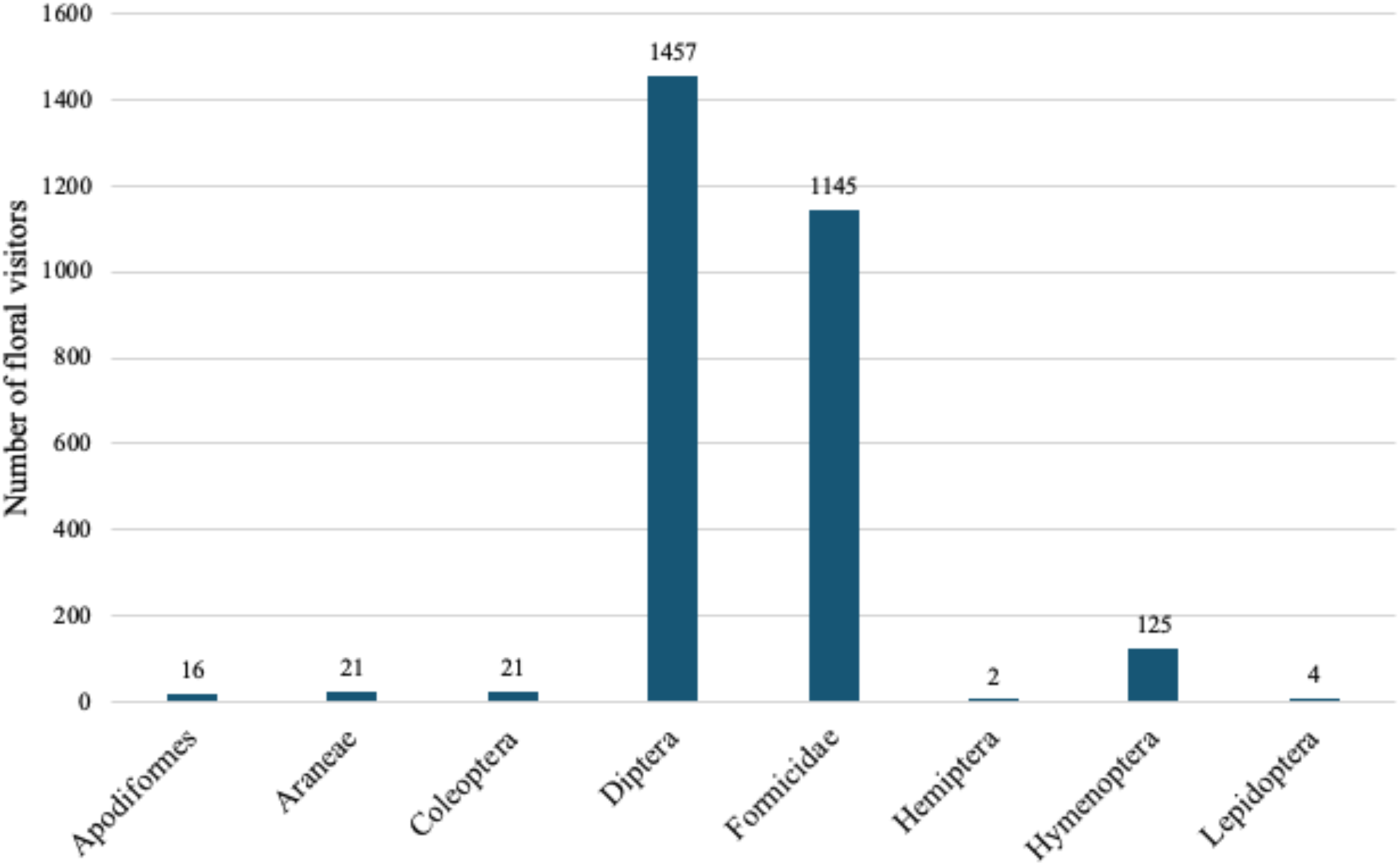
Summary of all artificial flower visitors. The horizontal axis labels correspond to taxonomic order of visitors, except for family Formicidae which is separated from the order Hymenoptera.

Consistent with our predictions, artificial flowers that were above average in size for each site attracted a greater number of floral visitors (*χ*² = 11.26, p < 0.001). There was also a significant effect of year (*χ*² = 46.25, p < 0.001), but the interaction of size and year, the effect of elevation, the quadratic effect of flower size were not significant (Fig. 6). This pattern held true when the dataset of floral visitors was reduced to only visitations by bees, butterflies, or hummingbirds (Size difference: *χ*² = 6.79, p = 0.009, Figure S4), which are considered effective pollinators across systems (Page et al., 2021).

**Figure 6:**
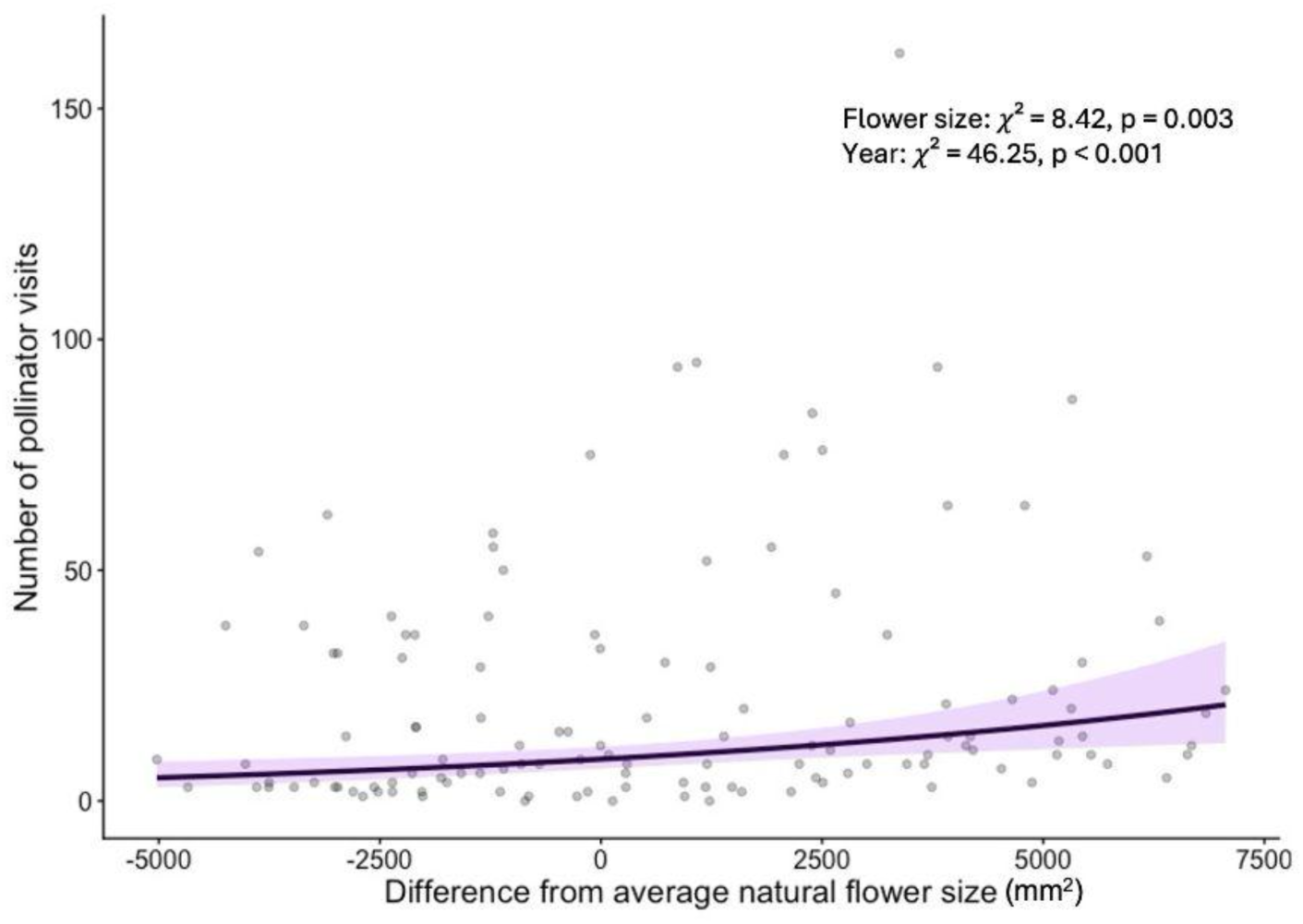
Association between the number of floral visitors and flower size. Flowers that are larger than average for the site receive more floral visitations. Each dot represents one floral visitation to an artificial flower. Purple line is a regression line from a generalized linear mixed model with a negative binomial distribution.

## Discussion

We found that *Iris missouriensis* flower size varies clinally along elevational gradients in the Rocky Mountains, but this pattern varies across years (Fig. 3), as the cline disappeared in 2023 when temperatures were significantly colder and precipitation was higher on average relative to other years of the study (Fig. 4). Thus, interannual variation in climate drivers could influence the expression of floral traits in this system dynamically across years such that cooler years with greater moisture availability relax stress induced reductions in floral size especially in the typically arid environments of low elevation locales. Additionally floral visitors are attracted to larger flowers consistently across the elevational gradient, even in arid locations where conditions likely induce the expression of smaller flowers (Fig. 6). This finding substantiates the hypothesis that the expression of flower size is likely driven by a trade-off between pollinator preference for larger flowers and the transpirational costs of larger flowers under arid conditions (Caruso et al., 2019; Day Briggs & Anderson, 2025; Siepielski et al., 2017; Sletvold, 2019; Strauss & Whittall, 2006). Collectively, these findings implicate fluctuations in abiotic conditions in shaping the variation in flower size across landscapes.

The clinal variation in flower size that we have detected in *Iris missouriensis* is consistent with patterns in other species of *Iris* (Lozada-Gobilard, Motter, et al., 2023; Sapir et al., 2002). For nine species of *Iris* native to the Middle East, flower size increases with latitude and increasing distance from the Negev desert (Sapir et al., 2002). Additionally, this pattern holds strong for a diversity of species outside of the Iridaceae naturally occurring across precipitation and aridity gradients (Lambrecht et al., 2017; Lambrecht & Dawson, 2007; Xie et al., 2016). For example, *Caragana stenophylla* (Fabaceae) produces smaller flowers in an intensively arid zone than a semi-arid zone (Xie et al., 2016). Additionally, water manipulation experiments in controlled chambers have generated similar clines in flower size in the species *Sinapis arvensis* (Brassicaceae) (Kuppler et al., 2021). Collectively, these studies, in combination with ours, suggest that aridity may drive clinal variation in flower size. Plasticity (Lambrecht et al., 2017) and genetic clines (Maad et al., 2013) could play a role in floral size, thus a common garden experiment is required to disentangle their relative roles in this system.

The preference of floral visitors for larger than average flowers was strong and consistent across sites (Fig. 6). Larger flowers often receive a greater number of floral visitations (Delgado et al., 2023; Kuppler et al., 2021; Thompson, 2001), and our study found that this pattern was maintained across an aridity gradient. Here, we exposed potential pollinators to artificial flowers that exceed the natural variation in floral size, allowing us to test if larger flowers receive more visitations than smaller flowers and if there is a limit to this preference. Since many bee and fly species have average foraging distances between 100-1000 meters (Rader et al., 2011; Zurbuchen et al., 2010), it is unlikely that individual pollinators regularly encounter populations with such drastically different average flower sizes highlighting an advantage of our experimental design. Floral visitors consistently preferred the largest available flowers, suggesting that size preference does not plateau and is not determined by the local average flower size. If the nutritional resources produced by these flowers also decline along with flower size under future conditions—as is expected under warming and drought scenarios (Kuppler & Kotowska, 2021; Moss & Evans, 2022)—pollinators could alter their foraging behavior (Phillips et al., 2018) or shift their ranges to more benign conditions (Day Briggs & Anderson, 2026; Gómez-Ruiz & Lacher Jr., 2019). We did detect an effect of year on the number of visitations to our artificial flowers indicating possible shifting pollinator communities abundances across years. This could be due to a variety of reasons that were unexplored in our study, but based on our analyses of climate data, 2021 was a warmer and dryer year relative to 2022, possibly resulting in fewer visitations in our 2021 dataset.

### Future directions

Significant post-flowering herbivory from ungulates prevented us from quantifying fecundity. Thus, we are unable to examine fecundity selection on flower size. We encourage future studies that focus on drought-mediated selection on floral size (Caruso et al., 2019; Day Briggs & Anderson, 2025), especially as climate change will increase aridity in this region (IPCC, 2023). Our findings suggest that the expression of smaller flowers could become more prevalent as climate change progresses, but the evolutionary consequences of this remains unclear. Larger bodies bees are more effective at depositing pollen, especially for larger flowers (Roquer-Beni et al., 2022; Willmer & Finlayson, 2014), but bee body size has declined over the past 100 years (Nooten & Rehan, 2020) due to increasing temperatures (Barrett et al., 2023). Given this observed shift, and reduced flower size under drought conditions, smaller-bodied bees could be more effective pollinators as climate change progresses.

*I. missouriensis* flowers are visited by potential pollinators capable of long-distance pollen dispersal (Day Briggs & Anderson, 2026). This was supported by our preliminary analysis of population structure that indicated spatially extensive gene flow. Thus, the clinal variation in floral structure that we have detected is likely not indicative of isolation-by-distance, but rather phenotypic plasticity or adaptation to spatially variable conditions. However, more extensive genetic analyses are required to confirm our results.

### Conclusions

We found that flower size of *Iris missouriensis* increases with elevation in the Colorado Rocky Mountains. Flower size is larger when there is greater precipitation that reduces the risk of excessive transpiration water loss. Climate was tightly correlated with elevation such that higher elevations are cooler and receive more precipitation. Through the use of 3D printed floral models, we found that potential pollinators prefer visiting flowers that are larger than average for a population. As climate change increases temperatures and drought conditions in this region, flowers will be reduced in size through phenotypic plasticity, but floral visitors will continue to prefer larger flowers.

