## Supplemental Information for "Pollinator preference for large flowers persists when climate induces the expression of smaller flowers"

| *Site number* | *Latitude* | *Longitude* | *Elevation (m)* | *Hours of natural flower recordings* | *Hours:Minutes of artificial flower recordings* |
|---|---|---|---|---|---|
| *1* | 39.106394 | -106.6037 | 3331 | 21 | 18:31 |
| *2* | 38.858439 | -107.09681 | 3010 | 24 | 21:16 |
| *3* | 38.973753 | -106.99618 | 2918 | 24 | 21:02 |
| *4* | 38.8948 | -107.0081 | 2723 | 21 | 22:14 |
| *5* | 38.822082 | -106.86909 | 2695 | 24 | 17:51 |
| *6* | 38.7301 | -106.8717 | 2527 | 15 | 20:39 |
| *7* | 38.53914 | -106.90602 | 2337 | 24 | 19:28 |

Supplemental Table S1: Table of recording site information. Site numbers correspond to the numbered blue circles in the site map of Figure 1A. Latitude, longitude, and elevation of each site are provided. The total number of hours of natural and artificial recordings at each site are also indicated.

| | Number of visits | |
|---|---|---|
| **Visitor ID** | Natural flower | Artificial flower |
| **Hyles lineata** | 0 | 1 |
| **Bombylius major** | 0 | 2 |
| **Protichneumon grandis** | 1 | 0 |
| **Selasphorus platycercus** | 4 | 16 |
| **Odonata sp.** | 1 | 0 |
| **Olla v-nigrum** | 1 | 1 |
| **Pristiphora sp.** | 1 | 0 |
| **Lassioglossum sp.** | 1 | 0 |
| **Urocerus flavicornis** | 1 | 0 |
| **Triepeolus sp.** | 1 | 0 |
| **Tharsalea rubidus** | 0 | 1 |
| **Papilio rutulus** | 6 | 0 |
| **Nomada sp.** | 14 | 5 |
| **Melissodes tristis** | 3 | 0 |
| **Neoclytus caprea** | 1 | 0 |
| **Polites draco** | 1 | 0 |
| **Megachilidae sp.** | 0 | 1 |
| **Hemiptera sp.** | 0 | 2 |
| **Hylaeus basalis** | 1 | 0 |
| **Hymenoptera sp.** | 10 | 1 |
| **Hoplitis sp.** | 0 | 1 |
| **Hoplitis fulgida** | 2 | 0 |
| **Hemaris thetis** | 3 | 1 |
| **Halictus rubicundus** | 1 | 0 |
| **Halictus tripartitus** | 4 | 1 |
| **Halictus sp.** | 16 | 1 |
| **Sphecodes sp.** | 0 | 1 |
| **Epeolus compactus** | 2 | 0 |
| **Halictidae sp.** | 3 | 4 |
| **Ebo evansae** | 1 | 0 |
| **Colletes sp.** | 7 | 1 |
| **Elasteridae sp.** | 16 | 0 |
| **Brephidium exilis** | 1 | 0 |
| **Chrysis angolensis** | 1 | 0 |
| **Coleoptera sp.** | 15 | 8 |
| **Nathalis iole** | 0 | 1 |

| Visitor ID | Natural flower | Artificial flower |
|---|---|---|
| **Muscidae sp.** | 27 | 209 |
| **Empididae sp.** | 5 | 7 |
| **Syrphidae sp.** | 20 | 107 |
| **Bombus insularis** | 4 | 0 |
| **Bombus grisecollis** | 2 | 0 |
| **Bombus flavidus** | 4 | 3 |
| **Bombus flavifrons** | 0 | 10 |
| **Bombus appositus** | 3 | 2 |
| **Bombus frigidus** | 1 | 0 |
| **Bombus morrisoni** | 1 | 4 |
| **Bombus huntii** | 3 | 13 |
| **Bombus kirbiellus** | 1 | 0 |
| **Bombus bifarius** | 1 | 2 |
| **Bombus melanopygus** | 3 | 0 |
| **Bombus californicus** | 2 | 1 |
| **Bombus nevadensis** | 2 | 0 |
| **Bombus sp.** | 1 | 1 |
| **Acmaeops proteus** | 1 | 0 |
| **Acrididae sp.** | 4 | 0 |
| **Andrena sp.** | 10 | 8 |
| **Agulla adnixa** | 1 | 0 |
| **Acmaeops pratensis** | 2 | 0 |
| **Agapostemon sp.** | 10 | 5 |
| **Osmia sp.** | 45 | 11 |
| **Ceratina sp.** | 60 | 46 |
| **Formicidae sp.** | 222 | 1151 |
| **Diptera sp.** | 432 | 1130 |
| **Andrenidae sp.** | 7 | 0 |
| **Apidae sp.** | 2 | 0 |
| **Arachnida sp.** | 10 | 0 |
| **Araneae sp.** | 1 | 21 |
| **Coccinellidae sp.** | 0 | 12 |

Supplemental Table S2: Identifications of floral visitors. Visitor ID is the finest grain identification that we were able to make for each visitor. The number of visitations are listed for natural flowers and artificial flowers.

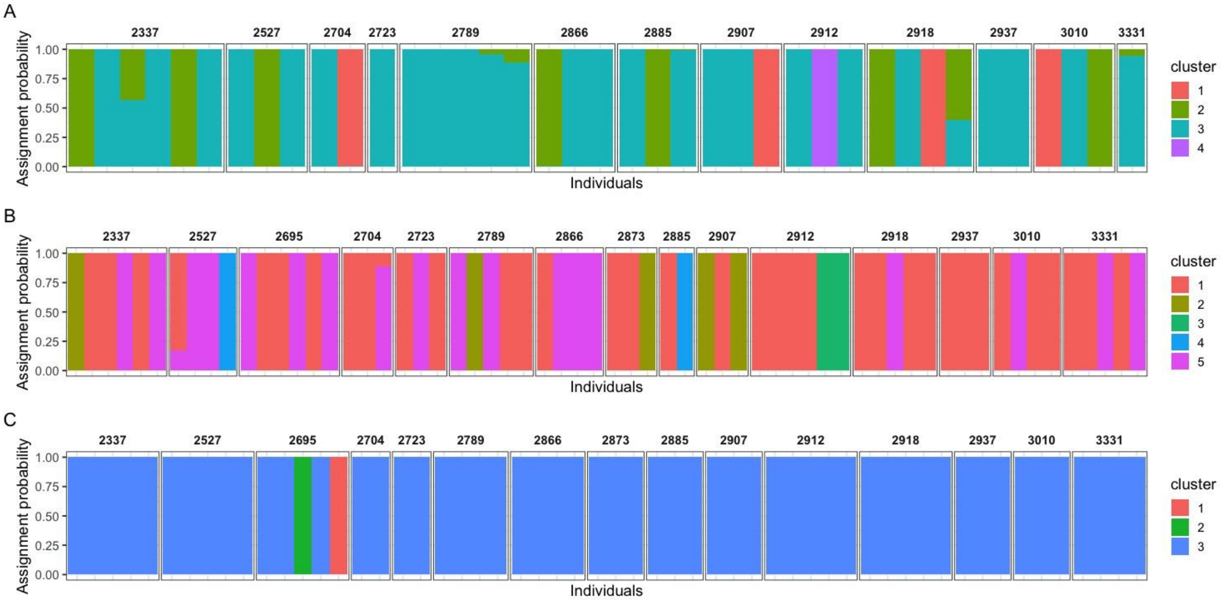


Supplemental Figure S1: Plots demonstrating population clustering across the 15 populations of Iris missouriensis for microsatellite markers A) IEST12, B) IEST24, and C)TrnL.

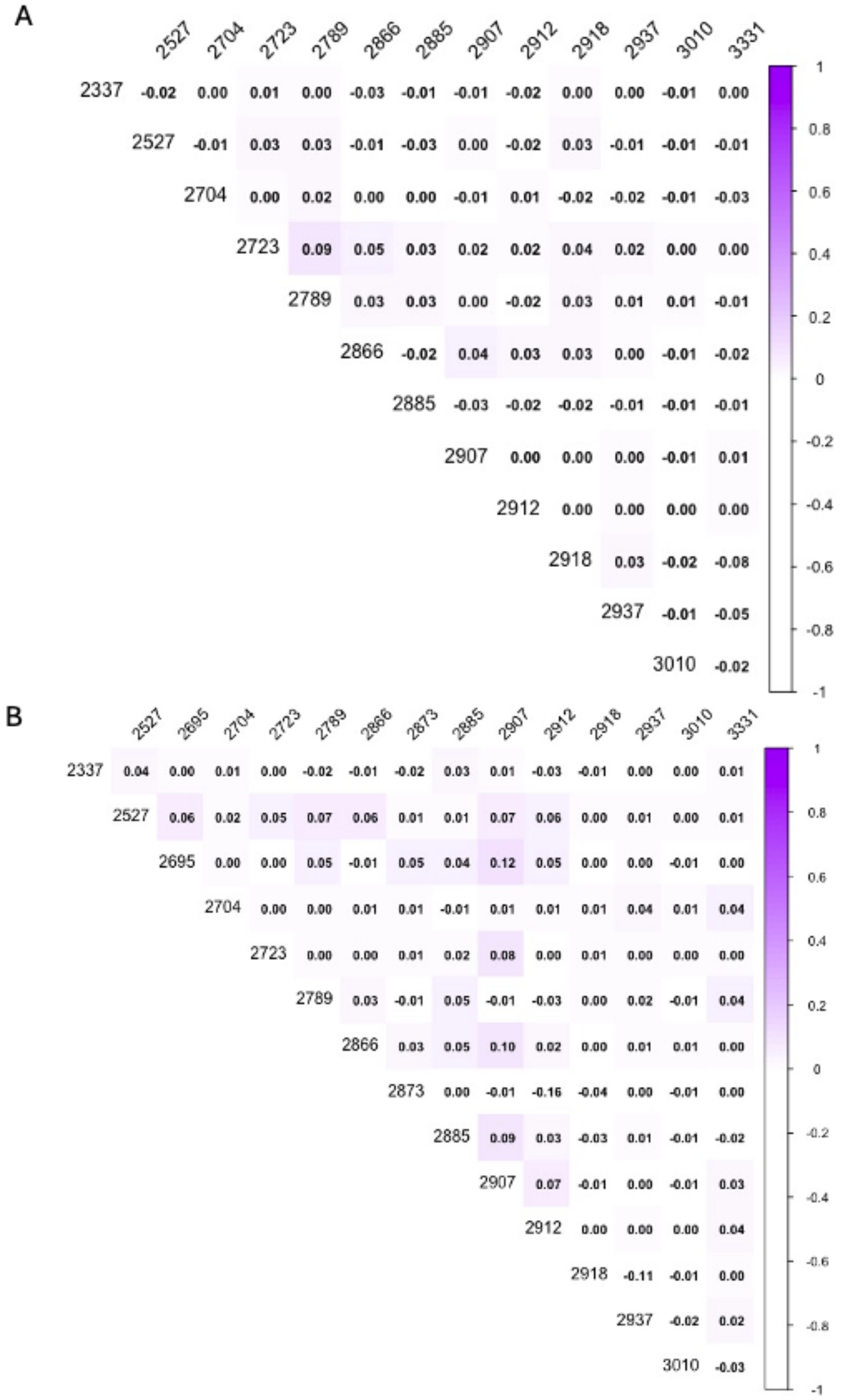


Supplemental Figure S2: Pairwise $F_{st}$ values calculated for the microsatellite A) IEST12 and B) IEST24.

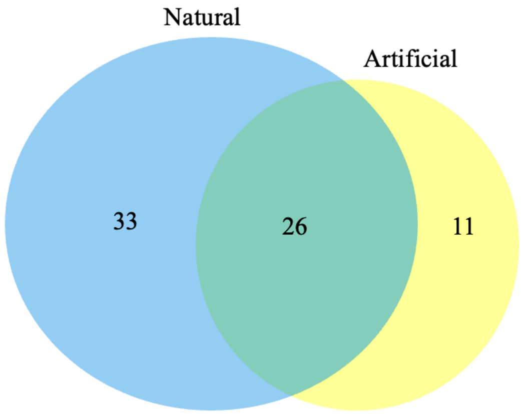


Supplemental Figure S3: Comparison of floral visitors to natural and artificial flowers. The numbers in the venn diagram indicate the number of unique species to visit each type of flower.

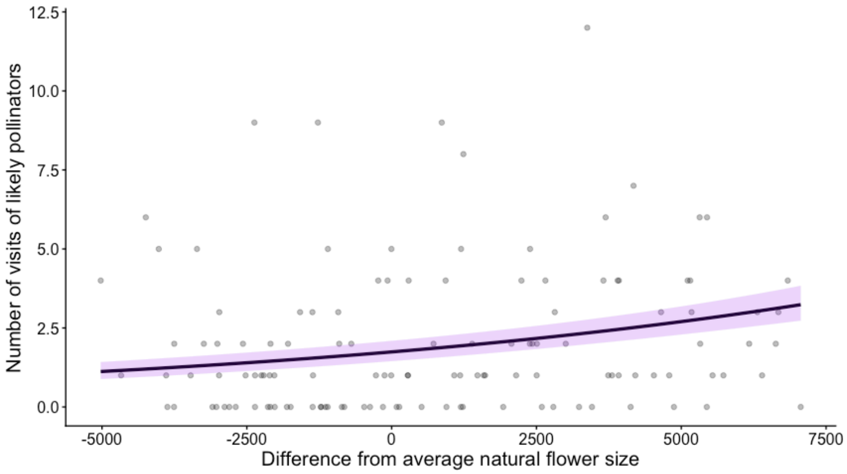


Supplemental Figure S4: The relationship between the number of visitations of “likely” pollinators and difference of flower size from population average. Larger than average flowers attracted a larger number of visitations of bee, butterfly, and hummingbird species.
